# A screen for sleep and starvation resistance identifies a wake-promoting role for the auxiliary channel unc79

**DOI:** 10.1101/2021.02.08.430360

**Authors:** Kazuma Murakami, Justin Palermo, Bethany A. Stanhope, Alex C. Keene

## Abstract

The regulation of sleep and metabolism are highly interconnected, and dysregulation of sleep is linked to metabolic diseases that include obesity, diabetes, and heart disease. Further, both acute and long-term changes in diet potently impact sleep duration and quality. To identify novel factors that modulate interactions between sleep and metabolic state, we performed a genetic screen for their roles in regulating sleep duration, starvation resistance, and starvation-dependent modulation of sleep. This screen identified a number of genes with potential roles in regulating sleep, metabolism or both processes. One such gene encodes the auxiliary ion channel UNC79, which was implicated in both the regulation of sleep and starvation resistance. Genetic knockdown or mutation of *unc79* results in flies with increased sleep duration, as well as increased starvation resistance. Previous findings have shown that *unc79* is required in pacemaker for 24-hour circadian rhythms. Here, we find that *unc79* functions in the mushroom body, but not pacemaker neurons, to regulate sleep duration and starvation resistance. Together, these findings reveal spatially localized separable functions of *unc79* in the regulation of circadian behavior, sleep, and metabolic function.

## Introduction

Sleep acutely regulates metabolic function, and growing evidence suggests that these processes interact to regulate many biological functions including cognition, physiology, and longevity (Hartmann 1974; Siegel 2005; Joiner 2016; Beckwith and French 2019). At the clinical level, many diseases related to metabolic dysfunction including diabetes, heart disease and obesity are associated with chronic sleep loss (Taheri *et al*. 2004; Arble *et al*. 2015; Reutrakul and Van Cauter 2018). In addition, diet potently influences sleep duration and quality, indicating that neural systems regulating sleep are sensitive to internal nutrient stores and food availability (Catterson *et al*. 2010; Grandner *et al*. 2010, 2014; Linford *et al*. 2012). Identifying how sleep, diet, and metabolic regulation are interconnected is critical to understanding the fundamental functions of sleep.

Interactions between sleep, feeding, and metabolic regulation are highly conserved between the fruit fly and mammals (Griffith 2013; Yurgel *et al*. 2015; Beckwith and French 2019). Experimental evolution and artificial selection approaches have revealed a relationship between sleep, feeding, and starvation resistance (Masek *et al*. 2014; Slocumb *et al*. 2015; Brown *et al*. 2019). For example, selection for short-sleeping flies results in reduced energy stores and sensitivity to starvation, while selecting for starvation resistance increases sleep duration (Seugnet *et al*. 2009; Masek *et al*. 2014; Slocumb *et al*. 2015). Further, examining naturally occurring genetic variation in sleep and starvation resistance in *Drosophila melanogaster* from different geographic localities suggests sleep and starvation resistance are inversely related (Brown *et al*. 2018; Sarikaya *et al*. 2020). The interactions between these traits under conditions of experimental evolution raise the possibility that shared genetic factors underlie sleep and starvation resistance.

Energy conservation has long been proposed to be primary function of sleep (Hartmann 1973; Berger and Phillips 1995). *Drosophila* live for only a few days in the absence of food, providing an excellent model to examine the effects of sleep on metabolic regulation and energy conservation (Yurgel *et al*. 2015; Ly *et al*. 2018). Quantifying longevity under starvation conditions provides a readout of overall energy stores and metabolic rate (Baldal *et al*. 2006; Schwasinger-Schmidt *et al*. 2012). In addition, flies acutely suppress sleep and increase activity in response to starvation, providing a system to investigate acute modulation of sleep and metabolic function (Lee and Park 2004; Keene *et al*. 2010). Genetic screens and genomic analyses have identified many regulators of sleep, metabolic regulation, and starvation resistance, establishing flies as a model for studying the interactions between these processes (Harbison *et al*. 2005; Jumbo-Lucioni *et al*. 2010; Murakami *et al*. 2016; Sonn *et al*. 2018). Many of the genes initially identified through screening for short-sleeping mutants have reduced life spans or increased sensitivity to stressors, though the relationship with starvation resistance is less clear (Koh *et al*. 2008; Bushey *et al*. 2010; Hill *et al*. 2018). A complete understanding of how these processes are integrated requires the localization of genes and neurons that regulate sleep and metabolic processes.

The study of sleep in flies has predominantly focused on the role of genes and neurons under fed conditions, leading to the identification of many distinct circuits that promote sleep and wakefulness (Allada and Siegel 2008; Sehgal and Mignot 2011; Ly *et al*. 2018). There is growing evidence that additional cell types are critical regulators of sleep including multiple classes of glia, endocrine cells, and the fat body (A rtiushin *et al*. 2018; Stahl *et al*. 2018; Vanderheyden *et al*. 2018; Yurgel *et al*. 2018; Ertekin *et al*. 2020). Further, the genes and neurons regulating sleep can differ based on environmental context (Griffith 2013; Beckwith and French 2019; Shafer and Keene 2021). These studies highlight brain-periphery interactions that are change in response numerous environmental contexts including food availability. Therefore, identifying genetic regulators that impact both sleep and metabolic function requires investigating both neuronal and non-neuronal cell-types.

Here, we have performed a genetic screen to identify genetic regulators of sleep and metabolic function, targeting genes ubiquitously to identify factors that function both within the brain and the periphery. Flies were tested in a pipeline that measured sleep parameters under fed and starved conditions, followed by assessment of starvation resistance. This screen identified several candidate genes regulating sleep and metabolic function including the sodium leak channel *NALCN* accessory subunit *unc79*. The phenotype of *unc79* is unique because it contributes to all three of these processes, suggesting *unc79* modulates both sleep and starvation resistance and potentially has a dual role in the regulation of sleep and metabolic function.

## Methods

### Fly husbandry

Flies for behavioral experiments were maintained and tested in humidified incubators at 25°C and under 65% humidity (Powers Scientific). Flies were reared on a 12 h:12 h light–dark cycle for experiments prior to behavioral analysis. All flies were maintained on Nutri-fly Drosophila food (Genesee Scientific)._All RNAi lines tested were obtained from the Bloomington *Drosophila* Stock Center (Bloomington, IN, USA) (Ni *et al*. 2008; Perkins *et al*. 2015) and the (# 45780) Vienna *Drosophila* Resource Center (Vienna, Austria) (Dietzl *et al*. 2007) (See table 1). Bridget Lear (Northwestern) generously provided *unc79* ^F01615^ and *unc79* ^F01615^ lines (Lear *et al*. 2013).

### Behavioral Analysis

*Drosophila* Activity Monitors (DAM; Trikinetics, Waltham, MA) were used for all behavioral analyses. The DAM system detects activity by monitoring infrared beam crossings for each animal (Pfeiffenberger *et al*. 2010a). These data were used to calculate sleep information by extracting immobility bouts of 5 minutes using the Drosophila Sleep Counting Macro (Pfeiffenberger *et al*. 2010b). All behavioral experiments used 5-7 day old mated female flies unless otherwise noted. For experiments examining the effects of starvation on sleep, activity was then measured for 24 hours on food, prior to transferring flies into tubes containing 1 % agar (Fisher Scientific) at ZT0 and activity was recorded for an additional 48 hours.

To measure starvation resistance, flies were starved on experimental day 2 by transferring them from food 1% agar (Fisher Scientific) individual DAM tubes containing 1% agar. Activity across the first 48 hours on agar was used to measure starvation-induced sleep suppression as previously described (Masek *et al*. 2014). While previous studies measured starvation-induced sleep suppression on day 1 of starvation, we measured this process for multiple days (Keene *et al*. 2010). To measure starvation resistance, activity was then recorded until death. Death was manually determined as the last activity time point from the final recorded activity bout for each individual fly.

For experiments quantifying circadian rhythm analysis, locomotor activity under free-running conditions was measured using the DAM system as previously described (Chiu *et al*. 2010). Individual flies were housed in 10% sucrose DAM tubes instead of standard fly food to prevent larval development that interferes that the circadian assay. Five day old adult flies were entrained to light-dark (LD) 12 hour: 12 hour (12:12) cycles for three days, then transferred to constant darkness (DD) for 7-8 days. Locomotor activity data were analyzed using Clocklab software (ActiMetrics, Version 2.72). Individual periods were calculated from 7-8 days activity data during DD using chi-square periodogram. Rhythm strength was determined by Fast Fourier Transform (FFT) analysis as previously described (Chiu *et al*. 2010).

### Statistical Analysis

Statistical analyses were performed using InStat software (GraphPad Software 6.0). For analysis of sleep, we employed a one- or two-way ANOVA followed by a Tukey’s post hoc test. For starvation resistance, we applied Kaplan–Meier analysis by grouping each genotype.

## Results

To screen for novel regulators of sleep and starvation resistance we ubiquitously knocked down genes by expressing RNAi transgenes from the TRiP Collection under control of the *Actin5C-GAL4* driver (Perkins *et al*. 2015). To enrich for genes that may be involved in sleep or metabolic function, candidate genes were selected from a genome-wide analysis of polymorphisms and genomic markers of selection in flies selectively bred for starvation resistance (Hardy *et al*. 2018). Sleep was measured for 24 hours on standard food, after which, flies were transferred to agar where they were maintained until death to measure starvation-induced sleep suppression and starvation resistance (Fig. 1A). In control flies harboring *Actin5C-GAL4* driving *UAS-luciferase-RNAi (Act5c>Luc^RNAi^*), a control with no endogenous targets, flies suppressed sleep during the first day of starvation, and an even greater suppression was observed on day two of starvation. Flies survived approximately three days on agar, providing a robust readout of sleep and starvation resistance.

**Figure 1.**
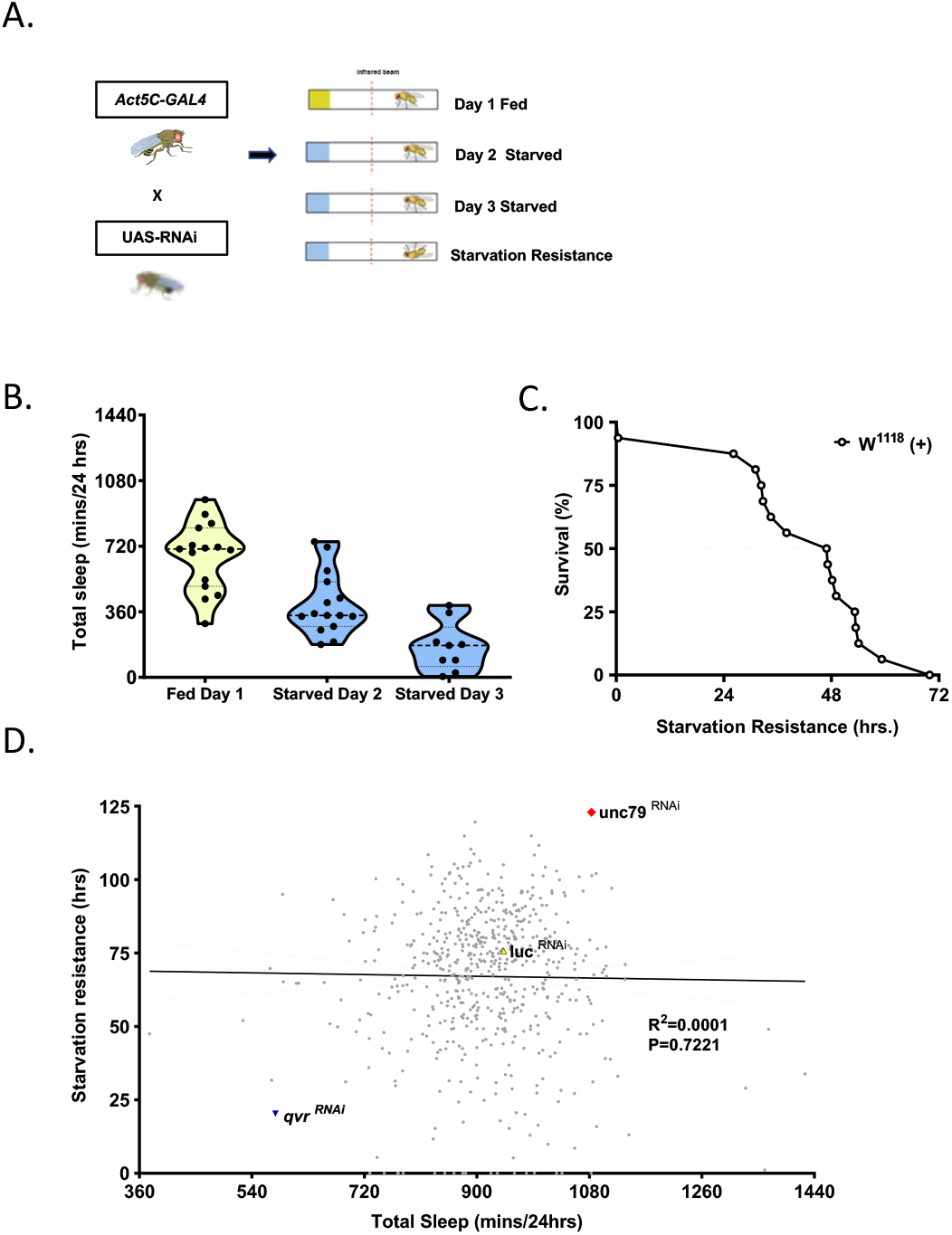
Screening for sleep and starvation resistance. **A.** Schematic of TRiP Screen. Ubiquitous Act5c GAL4 driver is crossed to TRiP lines, F1 flies were transferred to *Drosophila* Activity Monitor tubes. Sleep was measured for 24 hours on standard food, after which, flies were transferred to agar where they remained until death to quantify starvation-induced sleep suppression and starvation resistance. **B**. Flies suppressed sleep during the first day of starvation, and an even greater suppression was observed on day two of starvation. **C**. Flies survived approximately three days on agar, providing a robust readout sleep and starvation resistance. Total sleep (mins) is measured over 24 hour period and starvation resistance is measured in hours. **D.** Scatter plot for fed total sleep on x-axis plotted (mins) to starvation resistance on y-axis (hrs) (Simple Linear Regression: F (1,609) equals 0.1265, R^2^ value equal 0.0001, P-value > 0.7223). Control flies with no endogenous targets Act5c GAL4 drive *luc*^RNAi^ (yellow), lines tested in (grey), *qvr* has lowest sleep and SR (blue), and unc79 highest sleep and SR (red).

To identify novel regulators of these traits we sought to screen genes that were previously identified in a *Drosophila* genome-wide association study as regulating starvation resistance. Of the 1429 significant genes from this analysis, we identified 914 genes that TRiP RNAi stocks available (Perkins *et al*. 2015; Hardy *et al*. 2018). Of these, 299 lines (32.7%) were lethal with ubiquitous knockdown and, therefore, were not screened. In total, we screened 616 lines for sleep, starvation-induced sleep suppression (Fig. 1B and Table S1). Ubiquitous knockdown of the previously identified Ly-6 transmembrane protein, *qvr/sleepless*, resulted in the shortest sleep duration, confirming the ability of the screening procedure to effectively identify genetic regulators of sleep (Koh *et al*. 2008).

To examine the relationship between sleep and starvation resistance we plotted the average for each trait. There was no association between the traits (r^2^<0.001), suggesting these two traits are largely independently regulated (Fig. 1C). However, we identified a number of genes where ubiquitous knockdown resulted in increased sleep on food and greater starvation resistance (Fig. 1C). We also examined the correlation between genes screened for different sleep parameters. For example, daytime sleep duration is correlated with nighttime sleep duration, suggesting shared genes regulate both processes (Fig S1A). We found average bout length was inversely correlated with sleep bout number, suggesting these traits are functionally related (Fig. S1B). However, no correlation was observed between waking activity and total sleep (Fig. S1C) suggesting independent regulation of these traits. We chose to focus on the gene encoding for the NALCN auxiliary protein, *uncoordinated 79 (unc79*), because of the robustness of each phenotype and its role as an essential regulator of circadian rhythms sleep regulation (Lear *et al*. 2005; Joiner *et al*. 2013).

To validate the sleep and starvation phenotypes associated with *unc79* we repeated experiments and examined the sleep profile. Flies with ubiquitous knockdown of *unc79* (*Act5c*>*unc79*^RNAi^) slept significantly more than control flies (Fig. 2A,B). Further, while both control groups suppressed sleep during day 1 and 2 of starvation, *Act5c*>*unc79*^RNAi^ flies did not suppress sleep, suggesting that *unc79* is required for metabolic regulation of sleep (Fig. 2A,B). In addition, starvation resistance was significantly increased in *Act5c*>unc79^RNAi^ flies compared to *Act5c*>*Luc*^RNAi^ controls (Fig 2C). In agreement with our previous findings, waking activity in female control flies increased starvation, but was unchanged in *Act5C>Unc79^RNAi^* flies (Fig S2A) (Keene *et al*. 2010). Further, the overall waking activity was elevated in *Act5c*>*Unc79*^RNAi^ flies compared to *Act5c*>*Luc*^RNAi^ controls under fed conditions suggestions that the increased sleep in flies deficient for *unc79* is not due to general lethargy.

**Figure 2.**
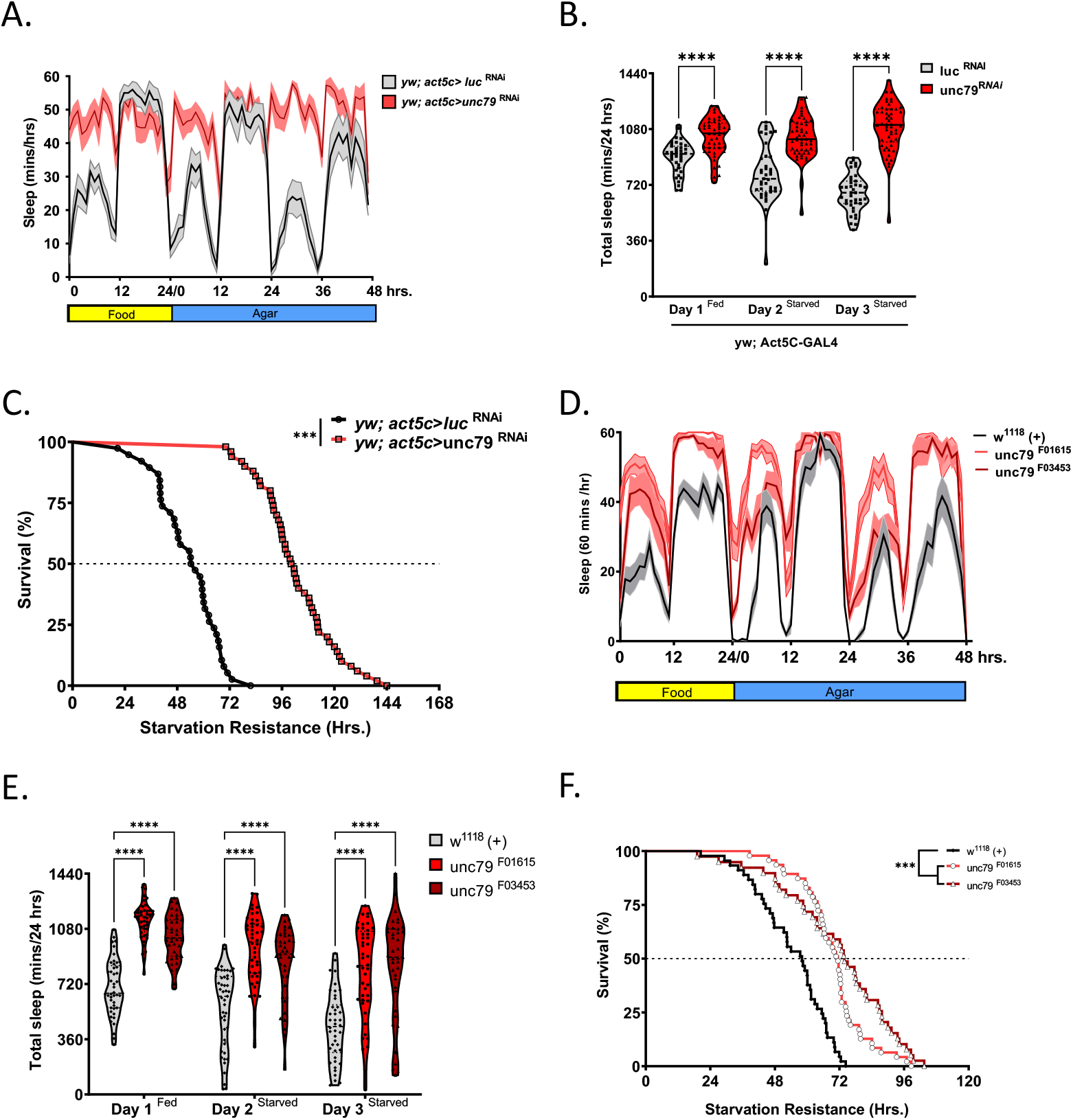
unc79 RNAi and mutants have increased sleep and starvation resistance. **A.** Sleep profiles depicting the average sleep each hour over a 72 hour experiment for Act5c>*luc*^RNAi^ (grey) and Act5c>*unc79*^RNAi^ (red). Flies were on food for day 1, then transferred to agar for days 2 and 3. **B.** Act5c>unc79^RNAi^ (red) during fed (Two-way ANOVA: F (2, 261) = 8.551, P<0.0001 N>39), starved day 1 (Two-way ANOVA: F (2, 261) = 8.551, P<0.0001, N>39) and starved day 2 (Two-way ANOVA: F (2, 261) = 8.551, P<0.0001 N>39) flies slept significatly longer compared to Act5c>luc ^RNAi^ (grey) controls. **C.** Starvation resistance of *Act5c>unc79^RNAi^* (red) is significatly higher than *Act5c>luc^RNAi^* (black) control (Gehan-Breslow-Wilcoxon test: Chi ^2^ = 94.42, df = 1, P-value < 0.0001). **D.** Sleep profile for hourly sleep averages over a 72 hour experiment for w^1118^, *unc79* ^F01615^ (red) and *unc79*^F03453^ (maroon) flies are on food for day 1, then transferred to agar for day 2 and 3. **E.** Total sleep is greater in *unc79*^F03453^ mutant under fed (maroon, F(2, 651) = 71.46, P<0.0001, N≥39), starved day 1 (P<0.001; N≥39) and starved day 2 (P<0.001; N≥39) conditions compared to control (grey). Total sleep is greater in *unc79*^F01615^ mutant (red) under fed (F(2, 651) = 71.46, P<0.0001, N≥39), starved day 1 (P<0.001; N≥39) and starved day 2 (P<0.001; N≥39) conditions compared to *w*^1118^ control (grey). **F.** Starvation resistance is greater in *unc79*^F03453^ (maroon, Gehan-Breslow-Wilcoxon test: Chi 2 = 29.1, df = 1, P-value < 0.0001) and *unc79*^F01615^ (red, (Gehan-Breslow-Wilcoxon test: Chi ^2^ = 18.6, df = 1, P-value < 0.0001) flied compared to w^1118^ control (grey). All sleep data are violin plots and SR data are survival curves. ****p < 0.0001.

To confirm that the observed phenotypes are not due to RNAi off targets we tested flies with a genetic mutation in the *unc79* locus. The independent Pbac element insertions in the *unc79* locus slept longer on food and failed to suppress sleep when starved, phenocopying RNAi knockdown (Fig. 2D,E) (Lear *et al*. 2013). Further, *unc79* mutants (*unc79^F03453^* and *unc79^F01615^*) survived significantly longer on agar than respective controls (Fig. 2F). Analysis of flies heterozygous for the mutation revealed the long sleeping phenotype is semi-dominant (Fig. 2D-F).

We also assessed male flies to determine whether these phenotypes generalize across sexes. Male *unc79* mutants (*unc79^F03453^* and *unc79^F01615^*) flies slept longer on food and failed to suppress sleep when starved (Fig S2A) and survived longer on agar than respective controls (Fig S2B), but the response was attenuated compared to female flies. Waking activity in male flies did not differ between fed and starved groups of *unc79* mutants, while controls increase waking activity (Fig S2C). Therefore, ubiquitous RNAi knockdown or genetic mutation of *unc79* results in increased sleep and starvation resistance, and impaired metabolic regulation of sleep.

To localize *unc79* function in metabolism and sleep we first targeted *unc79* ^RNAi^ to all neurons using the driver n-synaptobrevin-GAL4 (*nsyb*-GAL4) (Riabinina *et al*. 2015). Knockdown in neurons led to flies that slept significantly more than background controls harboring expressing RNAi to luciferase under fed conditions (Fig. 3A). Further, flies with pan-neuronal knockdown of *unc79* (*nSyb*-GAL4>*unc79*^RNAi^) also did not significantly reduce sleep during starvation and survived significant longer, suggesting *unc79* functions in neurons to regulate sleep and starvation resistance (Fig. 3A,B). To further localize the function of *unc79* we targeted RNAi to six types of neurons known to modulate sleep. Knockdown in the circadian neurons using *Pdf-GAL4* or *Tim-GAL4* did not affect sleep or starvation resistance, suggesting the effects on sleep and metabolic function are independent of its role in circadian activity (Fig. S3A). Further no effect was observed knocking down *unc79* in the sleep promoting central complex (*23E10-GAL4*) or broad classes of peptidergic cells (*C929*-GAL4) (Fig. S3A-D). Knockdown of *unc79* selectively in the mushroom bodies (*OK107-GAL4*) increased total sleep, specifically during the day, and resulted in increased starvation resistance (Fig. 3C-F). Therefore, selective knockdown with *OK107*-GAL4 largely phenocopies ubiquitous knockdown, raising the possibility that *unc79* functions in the mushroom body to regulate sleep and starvation resistance.

**Figure 3.**
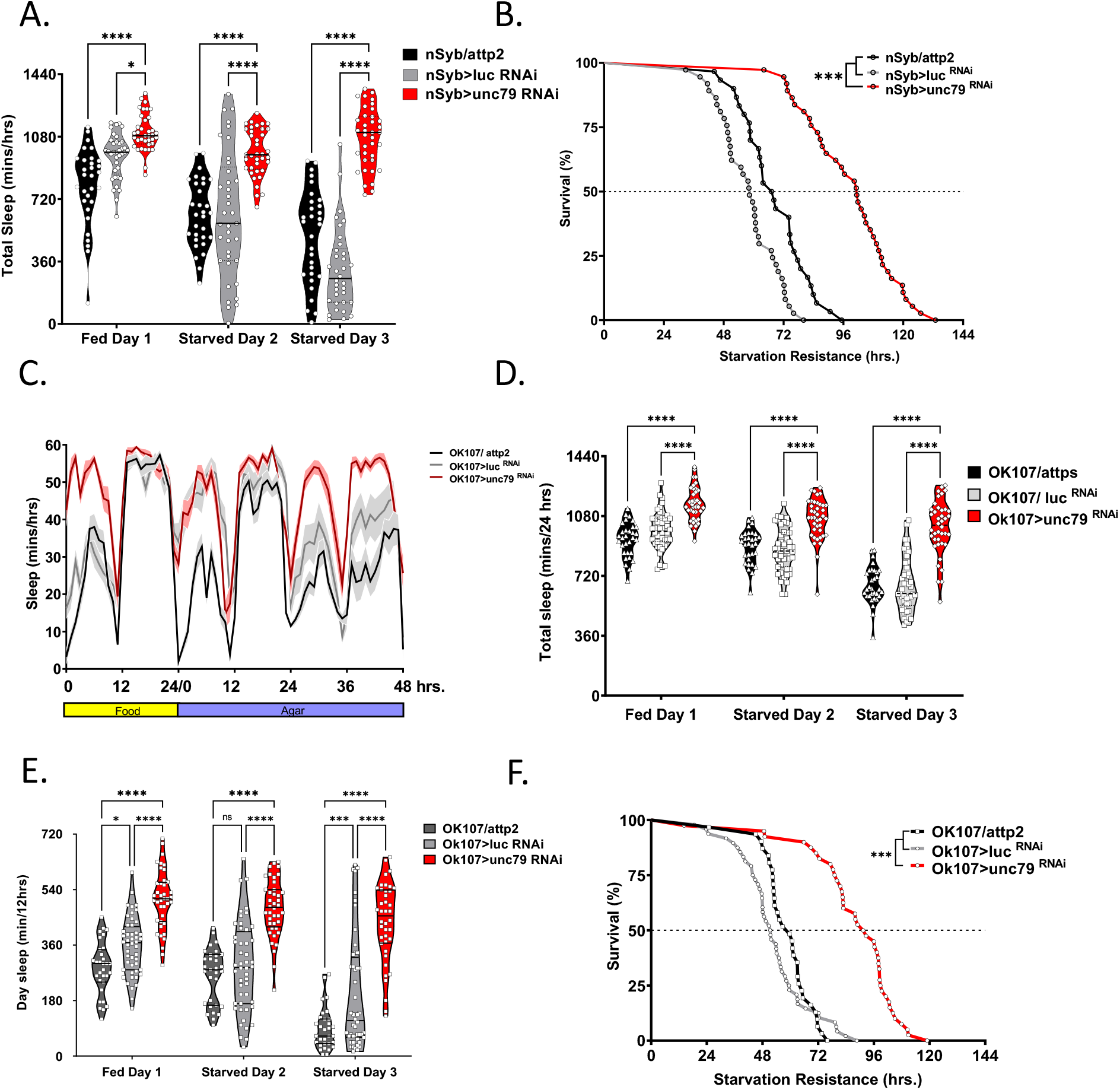
Localization of sleep and starvation resistance to the mushroom body. **A.** Pan-neuronal knockdown of *unc79* (nSyb>*unc79*^RNAi^, red) is significantly increased in sleep during fed (Two-Way ANOVA: F (2, 300) = 57.82, P<0.0001, N>30), starved day 1 (P<0.0001, N>30), and starved day 2 (P<0.0001, N>30); while, nSyb>attp2 (light grey) and nSyb>luc^RNAi^ controls shows starvation-induced sleep suppression. **B.** Starvation resistance for pan-neuronal knockdown of unc79 nSyb>*unc79*^RNAi^ (red) is significantly increased compared to nSyb>attp2 (grey, Gehan-Breslow-Wilcoxon test: Chi ^2^ equal 42, df equals 1, P–value<0.0001, N>30) and nSyb>luc^RNAi^ (light grey, Gehan-Breslow-Wilcoxon test: Chi ^2^ equals 64.6, df equals 1, P-value <0.0001, N>37) control flies. **C.** Sleep profile hourly sleep averages over a 72-hour experiment for mushroom body knockdown of unc79. Flies are on food for day 1, then transferred to agar for day 2 and 3. **D.** Mushroom body knockdown of unc79 (OK107>unc79 ^RNAi^, red) is significantly increased in sleep during fed (Two-Way ANOVA: F (2, 300) = 57.82, P<0.0001, N>30) starved day 1 (P<0.0001, N>31), and starved day 2 (P<0.0001, N>31); while, Ok107>attp2 (light grey) and Ok107>*luc*^RNAi^ controls shows starvation-induced sleep suppression. **E.** Daytime sleep in flies with mushroom body knockdown of unc79 (OK107>unc79 RNAi, red) is significantly increased under fed conditions (Two-Way ANOVA: F (2, 348) = 43.42, P<0.0001, N>30), starved day 1 (P<0.0001, N>31), and starved day 2 (P<0.0001, N>31); while, nSyb>attp2 (grey) and nSyb>*luc*^RNAi^ (light grey) controls maintain normal daytime sleep. **F.** Starvation resistance is increased in OK107>*unc79*^RNAi^ flies compared to Ok107>attp2 (grey, Gehan-Breslow-Wilcoxon test: Chi 2 equals 47.6, df equals 1, P-value <0.0001, N>31) and OK107>*luc*^RNAi^ (light grey, Gehan-Breslow-Wilcoxon test: Chi ^2^ equals 50.5, df equals 1, P-value <0.0001, N>31) control flies. All sleep data represent violin plots and SR data are survival curves. ****p < 0.0001.

The driver *OK107*-GAL4 expresses in some neurons that are extrinsic to the mushroom bodies including the Pars Intercerebralis (Aso *et al*. 2009). To determine whether the phenotypes observed localize to the mushroom body we expressed *unc79*^RNAi^ using *R13F02-GAL4*, a highly selective driver for the mushroom body (Jenett *et al*. 2012). These flies also slept longer than controls, failed to suppress sleep and survived longer under starvation conditions (Fig. 3G-H). Similar to findings with *OK107-GAL4*, knockdown of *unc79* in the mushroom bodies body with *R13F02* increased daytime, but not nighttime sleep (Fig. S3E-F). Therefore, these findings confirm that *unc79* functions in the mushroom bodies to regulate sleep and starvation resistance.

To determine whether the phenotypes observed are specific lobes of the mushroom body, we tested the effects of *unc79* knockdown in the α/β lobes (*c739*-GAL4), α’/β’ lobes (*c305a*-GAL4), and the γ lobes (*1471* GAL4 drivers) (Krashes *et al*. 2007; Aso *et al*. 2009). Flies with *unc79* knockdown in α/β lobes and α’/β’ lobes fail to suppress sleep compared to *luc* control, while knockdown in the γ lobes increased total sleep duration compared to *luc* control and fail to suppress sleep when starved (Fig. 4A). Starvation resistance is increased in *unc79* knockdown in each mushroom body subtype compared to their respective controls expressing *luciferase-RNAi* (Fig. 4B). Therefore, loss of *unc79* function in each subset of mushroom body neurons impacts sleep and metabolic regulation, while selective loss in the γ lobes largely recapitulates the full extent of ubiquitous knockdown. These findings reveal that *unc79* is required in all lobes of the mushroom body for proper sleep and metabolic regulation.

**Figure 4.**
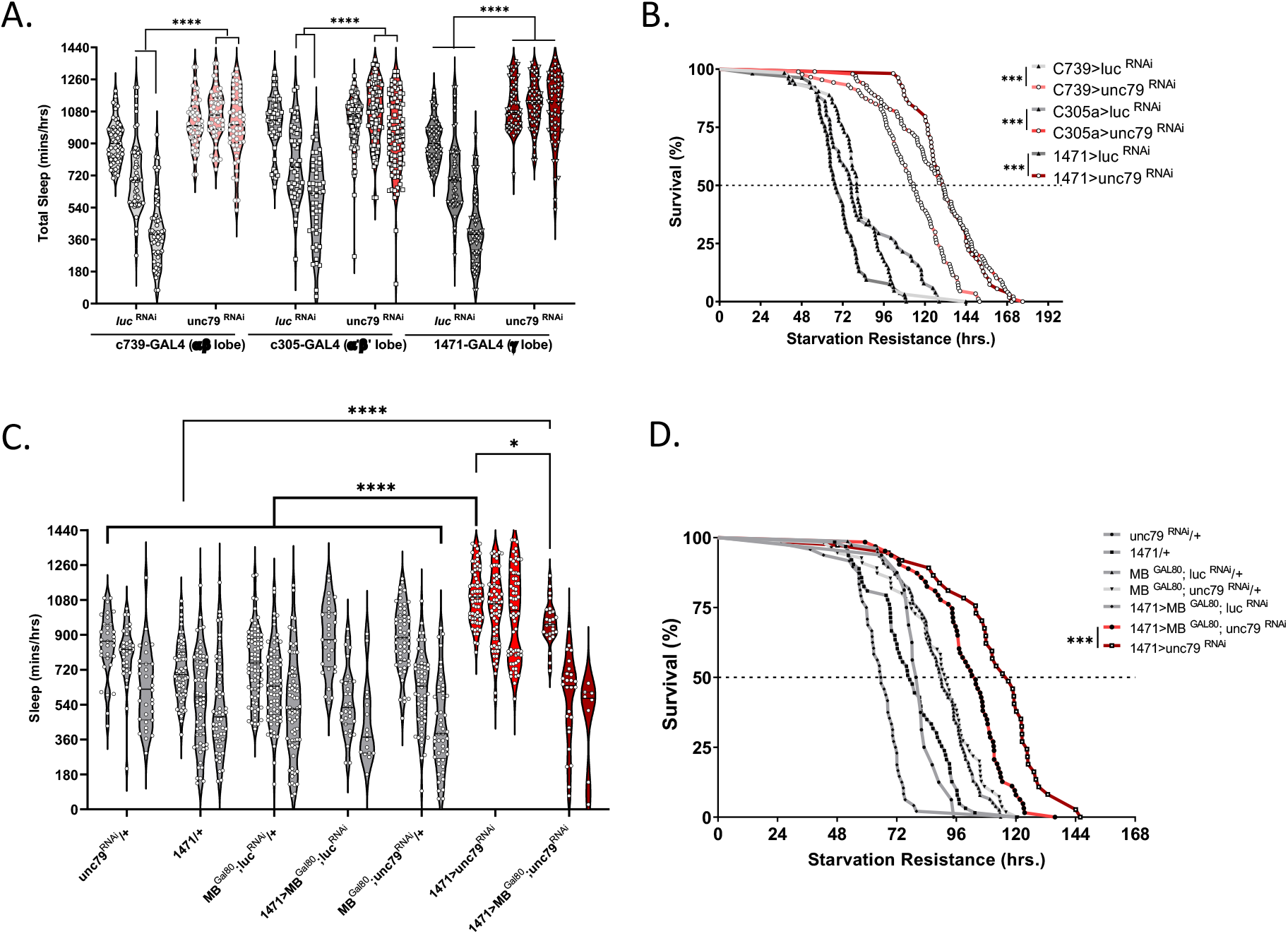
*unc79* function in the in mushroom body γ-lobe to regulate sleep and starvation resistance. **A.** Flies with *unc79* knocked down in the αβ lobes (c739>unc79 RNAi, pink) fail to suppress starvation during starved day 1 (Two-way ANOVA F (10, 1075) = 28.56, P<0.0001, N>53) and starved day 2 (P<0,0001) compared to control (c739>luc ^RNAi^, light grey); while fed day 1 did not differ (P>0.1587). Mushroom body α’β’ knockdown of unc79 (c305a> unc79 RNAi, red) fails to suppress starvation during starved day 1 (Two-way ANOVA F (10, 1075) = 28.56, P<0.0001, N>51) and starved day 2 (P<0.0001) compared to control (c305a>luc RNAi, light grey); while fed day 1 did not differ (P>0.999). Mushroom body γ knockdown of unc79 (1471 >unc79^RNAi^, maroon) significantly increase total sleep during fed day (Two-way ANOVA F (10, 1075) = 28.56, P<0.0001, N>53) and fails to suppress sleep on starved day 1(P< 0.0001), and starved day 2 (P<0.0001) compared to control (1471>luc^RNAi^, dark grey). **B.** Starvation resistance increased when mushroom body αβ knockdown of unc79 (c739 >unc79 ^RNAi^, pink) is significant (Gehan-Breslow-Wilcoxon test: Chi ^2^ equals 71.36, df equals 1, P-value<0.0001, N>63) compared to control (c739>luc^RNAi^, light grey). Starvation resistance increased when mushroom body α’β’ knockdown of unc79 (c305a >unc79^RNAi^, red) is significant (Gehan-Breslow-Wilcoxon test: Chi ^2^ equals 90.45, Df equals 1, P-value <0.0001, N>51) compared to control (c305a>luc ^RNAi^, grey). Mushroom body γ knockdown of *unc79* (1471 >unc79^RNAi^, maroon) is significant (Gehan-Breslow-Wilcoxon test: Chi ^2^ equals 124.6, df equals 1, P-value<0.0001, N>53) compared to control (1471>luc RNAi, dark grey). **C.** Mushroom body GAL80 rescue γ knockdown of unc79^RNAi^ (1471>MB^Gal80^; *unc79*^RNAi^) significantly rescues total sleep compared to γ knockdown of unc79 RNAi (1471>*unc79*^RNAi^, P-value <0.0432); while, total sleep for γ knockdown of *unc79*^RNAi^ (1471>*unc79*^RNAi^) is high compares to other control groups (MBGal80;lucRNAi/+, P-value <0.0001; *unc79*^RNAi^/+, P-value <0.0001; 1471/+, P-value <0.0001; MBGal80;*luc*-/+, P-value <0.0001; 1471>MBGal80;*luc*^RNAi^, P-value <0.0001; and, MBGal80;*unc79*^RNAi^/+, P-value <0.0001). Mushroom body GAL80 rescue γ knockdown with *unc79*^RNAi^ (1471>MBGal80; unc79 RNAi) restored total sleep to controls (*unc79*^RNAi^/+, P-value equal 0.3715; MBGal80;*unc79*^RNAi^/+, P-value equals 0.7292; and, 1471>MBGal80;*luc*^RNAi^, P-value quals 0.8177). However, 1471>MBGal80;*unc79*^RNAi^ vs.1471/+ remained significant (P–value<0.0001). **D.** Starvation resistance of mushroom body GAL80 rescue γ knockdown of unc79 RNAi (1471>MBGal80; *unc79*^RNAi^) significantly lower (Gehan-Breslow-Wilcoxon test: Chi ^2^ equals 11.13, Df equals 1, P-value<0.0009, N>37) compared to mushroom body γ knockdown of *unc79* (1471 >*unc79*^RNAi^, maroon). All sleep data are violin plots and SR data are survival curves. ***p < 0.001; ****p < 0.0001.

To verify that the increase in sleep and starvation resistance is specific to the mushroom bodies, we examined whether including of the *MB*-GAL80 transgene reverses the effects of *unc79* knockdown. Expression of GAL80 in the mushroom bodies restores sleep and starvation-induced sleep suppression to control levels (Fig. 4C). Starvation resistance increased in flies with *unc79* selectively knocked down in the γ lobes (*1471-* GAL4>*unc79*^RNAi^). Blocking expression of GAL4 within the mushroom body restored starvation resistance, and partially restored sleep duration on food, confirming that loss of *unc79* in the mushroom body leads to dysregulated sleep (Fig. 4C). In addition, the expression of *MB*-GAL80 restored normal starvation resistance to *1471*-GAL4>*unc79*^RNAi^ flies. Therefore, loss of *unc79* function within the γ-lobe of mushroom bodies increases both sleep and starvation resistance.

Previous work has revealed that *unc79* functions within the circadian neurons in association with the *unc80* accessory protein and the ion channel *narrow abdomen (na*) to maintain locomotor rhythms during constant darkness (Lear *et al*. 2005, 2013; Moose *et al*. 2017). To further investigate whether the sleep and circadian phenotypes are controlled by shared or distinct neural circuits, we knocked down additional components of the *unc79* complex in the mushroom bodies and measured the effects on sleep and starvation resistance. Knockdown of *na* or *unc80* throughout the mushroom bodies did not increase sleep duration on food or disrupt starvation-induced sleep suppression (Fig 5A). In addition, knockdown of *unc*80 and *na* in the mushroom body had little impact on starvation resistance (Fig 5B). These findings suggest *unc79* functions independently of its canonical complex with *unc80* and *na* to regulate sleep and starvation resistances.

**Figure 5.**
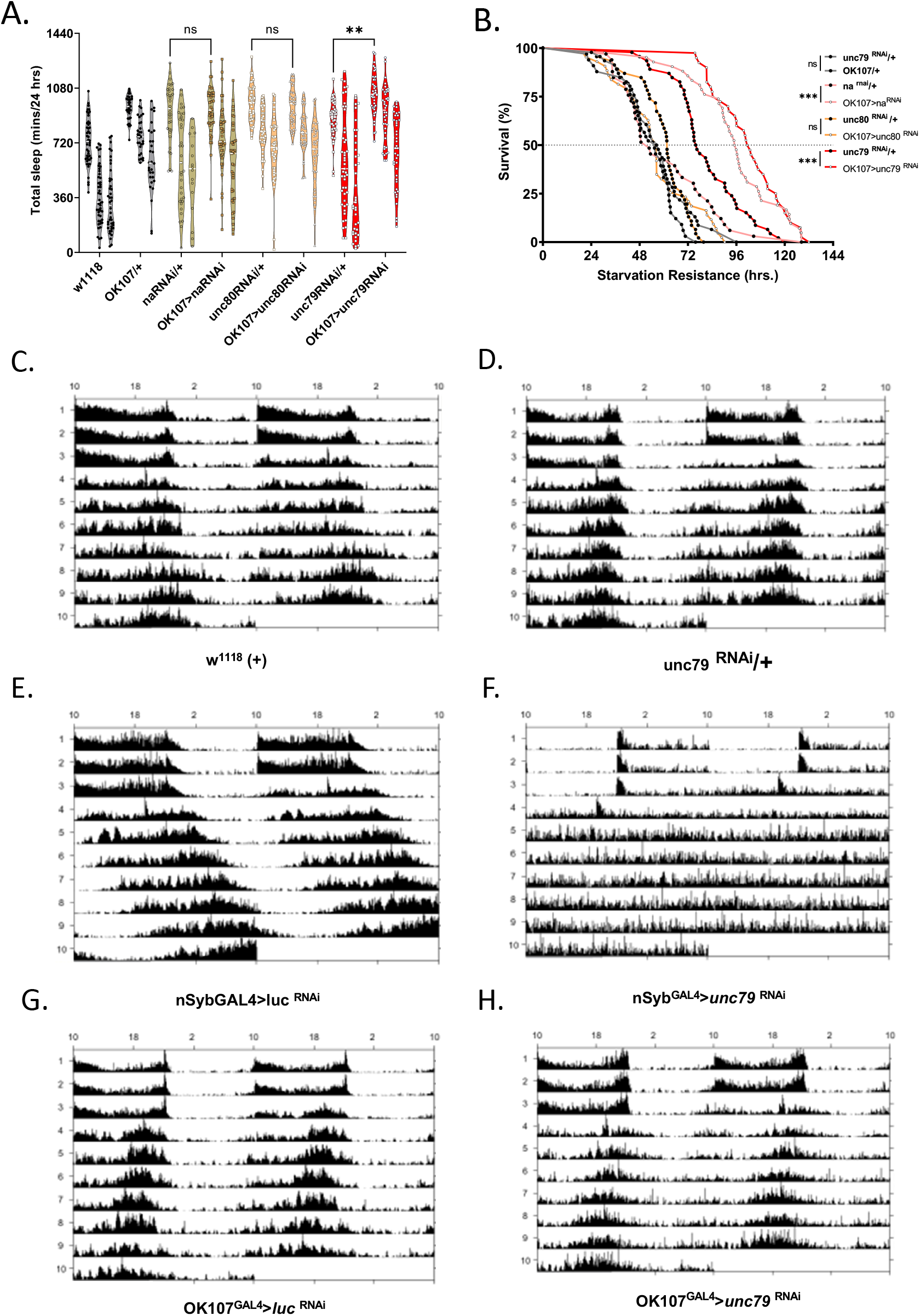
MB knockdown of unc79-RNAi does not impact circadian rhythms. **A.** Mushroom body knockdown of *unc79* (OK107>unc79RNAi) significantly increased sleep (Two-way ANOVA: F (2, 711) = 169, P<0.0029, N>40) compared to control (*unc79*^RNAi/+^). Mushroom body knockdown of narrow abdomen (OK107>*na*^RNAi^) did not differ (P>0.9999, N>35) compared to control (*na-*RNAi/+). Mushroom body knockdown of unc80 (OK107>unc80RNAi) did not differ (P>0.9964, N>42) compared to control (unc80 RNAi/+). **B**. Starvation resistance mushroom body knockdown of unc79 (OK107>unc79 RNAi) is increased (Gehan-Breslow-Wilcoxon test: Chi ^2^ equal 25.95, df equal 1, P-value <0.0001, N>40) compared to control (unc79-RNAi/+). Starvation resistance mushroom body knockdown of narrow abdomen (OK107>na^RNAi^) is increased (Gehan-Breslow-Wilcoxon test: Chi ^2^ equal 33.54, df equal 1, P-value <0.0001, N>42) compared to control (*na*^RNAi/+)^. Starvation resistance mushroom body knockdown of narrow abdomen (OK107>*unc80*^RNAi^) is no different (Gehan-Breslow-Wilcoxon test: Chi ^2^ equal 2.486, df equal 1, P-value <0.1148, N>43) compared to control (*unc80*^RNAi^/+). **C.-H.** Actogram double plot. Female flies were entrained in light-dark cycle for days 1-3 and dark-dark cycles 4-10 days. A, B, C, E and F have normal rhythm. Circadian rhythm is disrupted when unc79 is knocked down pan-neuronally, while unc79 knockdown in mushroom body has restored rhythm. All sleep data are violin plots and SR data are survival curves. ***p < 0.001; ****p < 0.0001.

Expression of *unc79* is required within pacemaker neurons to circadian rhythms. Therefore, it is possible that *unc79* functions in distinct populations of neurons to regulate circadian rhythms, from those regulating sleep and metabolic phenotypes. To directly test this possibility, we measured the effects of mushroom body-specific knockdown of *unc79* on free-running activity in entrained animals. As expected, flies with pan-neuronal knockdown are arrhythmic under conditions of constant darkness, while control flies show robust rhythms (Fig 5C-F). Conversely, knockdown in OK107-expressing cells does not impact circadian activity (Fig. S4; Fig 5G-H). Therefore, *unc79* function to regulate circadian rhythms through distinct neural mechanisms that regulate sleep and starvation resistance.

## Discussion

Here, we screened by targeting gene function ubiquitously to identify regulators of sleep and metabolic function. Growing evidence suggests sleep is regulated by complex interactions between the brain and periphery, including the findings that mutants impacting fat storage, and communication from the fat body to the brain significantly impact sleep (Thimgan *et al*. 2010; Slocumb *et al*. 2015; Ertekin *et al*. 2020).

We have identified numerous candidate regulators of sleep, including a novel role for *unc79* in the regulation of sleep and metabolic function. *Unc79* and *unc80* are auxiliary subunits of the sodium leak channel *na*, an ortholog of mammalian *NALCN* family of ion channels (Swayne *et al*. 2009). A number of functions have been identified for this complex including a role in the regulation of circadian rhythms, and anesthesia sensitivity (Lear *et al*. 2005; Humphrey *et al*. 2007). Previous work found that loss of narrow abdomen or *unc79* increased sensitivity to the anesthetics, halothane and isofluorane, and increases sleep (Humphrey *etal*. 2007; Joiner *etal*. 2013) consistent with our findings of increased quiescence in *unc79* mutants. Mutation of *unc79* also facilitates the emergence from anesthesia, raising the possibility that loss of *unc79* promotes state transitions, rather than directly impacting isofluorane sensitivity (Joiner *et al*. 2013). Therefore, suppression of arousal may be involved in anesthesia and sleep (Joiner *et al*. 2013). Mutation of *na* also impacts a number of complex behaviors including social clustering (the distance maintained between individual flies) (Burg *et al*. 2013), and light-mediated locomotor activity (Nash *et al*. 2002).These findings suggest a complex role for *na* and associated *unc79* genes in regulating brain function.

Multiple lines of evidence suggest the role of *unc79* in the regulation of sleep, metabolic regulation of sleep, and starvation resistance is separate from its essential role in regulating circadian rhythms. First, we localize function to the mushroom body, a region that is critical for regulation of sleep and modulation of behavior in accordance with feeding state (Joiner *et al*. 2006; Pitman *et al*. 2006; Sitaraman *et al*. 2015a; Tsao *et al*. 2018). We previously reported that the mushroom bodies are dispensable for starvation-induced sleep suppression, however the manipulations that led to this conclusion involved pharmacological ablation or acute genetic silencing of the mushroom bodies (Keene *et al*. 2010). Therefore, it is possible that loss of *unc79* function impacts sleep circuitry through a mechanism that would not be detected in flies with the previously applied genetic manipulations. Second, *na* and *unc80*, two components of a complex that interacts with *unc79* to regulate circadian rhythms, are dispensable for regulation of sleep and starvation resistance in the mushroom bodies. These findings raise the possibility that *unc79* may function independently of its canonical complex with *unc80* and *na*. A central question is how *unc79* functions to modulate mushroom body physiology and sleep circuitry. Studies examining the role of *unc79* in circadian function and anesthesia sensitivity suggests it functions by regulating *na* activity to modulate neural activity (Moose *et al*. 2017), and it is possible that *unc79* modulates the function of a different ion channel within the mushroom bodies.

We identify three independent phenotypes to the mushroom bodies. First, we find knockdown of *unc79* in the mushroom bodies promotes sleep suggesting a wake-promoting role for mushroom bodies. The mushroom bodies contain both wake and sleep-promoting neurons, and genetic ablation or silencing of the mushroom body increases wakefulness (Joiner *et al*. 2006; Pitman *et al*. 2006; Sitaraman *et al*. 2015a). It is possible that loss of unc79 is functioning in either sleep promoting or wake-promoting neurons to elicit this phenotype. We also identify two independent metabolic phenotypes to the mushroom bodies. In *Drosophila*, the mushroom circuits have been well-defined including the identification of modulatory neurons, and output neurons that modulate sleep (Aso *et al*. 2014; Haynes *et al*. 2015; Sitaraman *et al*. 2015b). Therefore, the identification of *unc79* as a regulator of sleep provides the opportunity to examine how gamma lobe output neurons are regulated by input neurons and impact the physiology of output neurons.

In addition to the sleep phenotypes, we find *unc79* mutants are resistant to starvation. This finding is particularly interesting because the list of genes chosen for the screen derived from those identified in a Genome Wide Analysis Study for factors associated with starvation resistance (Harbison *et al*. 2004; Hardy *et al*. 2018). Many different factors contribute to starvation resistance including energy stores, basal metabolic rate, and changes in metabolic rate upon starvation. Animals selected for starvation resistance have elevated sleep and do not suppress sleep when starved (Masek *et al*. 2014). Therefore, future work studying starvation selected lines, or other populations of outbred fly lines have potential to identify whether variable expression of *unc79* is associated with naturally occurring differences in sleep and metabolic regulation.

We find that *unc79* most potently impacts sleep and starvation resistance in the gamma lobes, suggesting this population is critical for both sleep metabolic regulation. Output neurons from the gamma lobes have been directly implicated in feeding and fat storage supporting the notion that this region is critical for metabolic regulation (Al-Anzi and Zinn 2018). Future work examining the effects of *unc79* deficiency on the physiology and function of mushroom body output neurons may help identify the role of *unc79* in regulating mushroom body circuits that ultimately regulate behavior and metabolic function. Taken together, these findings add to growing evidence that sleep and metabolic function are integrated. The identification of additional genetic factors that regulate the relationship between sleep and nutritional state through behavioral studies will improve our understanding of the strong associations between sleep loss and metabolism-related diseases. The ubiquitous screen has identified numerous candidate genes that impact sleep, starvation-induced sleep suppression, and starvation resistance, providing candidates that function within and outside of the nervous system. Future study of these genes, such as *unc79*, has potential to advance our understanding of sleep-metabolism interactions and brain-periphery communication.

## Acknowledgements

This work was supported by National Institute of Health awards R01HL143790 and R01DC01790 to ACK. The authors are grateful to Peter Lewis (FAU) for technical support and Allen Gibbs (UNLV) for helpful discussions and sharing unpublished data.

---

**Supplemental Figure 1.**

**A.** Scatter plot depicting day time and nighttime sleep scatter plot. Fed day sleep on x-axis plotted (mins/12 hrs) to Fed night sleep on y-axis (mins/12 hrs) (Simple Linear Regression: F (1, 609) equals 57.65, R^2^ equals 0.0863, P-value equals<0.0001). **B.** Scatter plot depicting the average number of sleep bouts compared to the average bout length. Sleep bout on x-axis to average bout length on y-axis Simple Linear Regression: F (1, 611) equals 338.2, R^2^ 0.3563, P value <0.0001). **C.** Total sleep/waking activity scatter plot. Fed total sleep on x-axis plotted (mins/24 hrs) to waking activity on y-axis (beam breaks/min (Simple Linear Regression: F (1, 612) equals 4.568, R^2^ equals 0.0074, P-value equals<0.0330). Control flies with no endogenous targets Act5c GAL4 drive luc^RNAi^ (yellow), lines tested in (grey), and unc79 highest sleep and SR (red).

**Supplemental Figure 2.**

**A.** Waking activity violin plots over a 72-hour experiment for female Act5c>lucRNAi (grey) and Act5c>unc79RNAi (red). Waking activity is higher during Day 1 fed (N≥37; P<0.0001) and Day 3 starved (N≥37; P<0.01), while not different during Day 2 (N≥32; P<.1199). **B.** Waking activity violin plots over a 72-hour experiment for female unc79 mutants. Waking activity differ during starved state of *unc79*^F03453^ (maroon) *unc79*^F01615^ (red) more than *w1118* (grey) and *unc79/+* controls (pink) (N≥39; P<0.0001), but not during fed state (N≥39; P<0.856). **C.** Total sleep violin plot over a 48-hour experiment for male *unc79* mutants. Waking activity differ during fed and starved state of unc79F01615 (red) more than *w1118* (grey) and *unc79/+* controls (pink) (N≥39; P<0.0001), but not during fed or starved state for *unc79*^F03453^ (maroon) (N≥39; P<0.8559). **D.** Starvation resistance of male flies did not differ between *unc79*^F03453^ (maroon) *unc79*^F01615^ flies (red), and both were greater than *w1118* (grey) and *unc79/+* controls (pink). E. Waking activity for male flies for 48-hour period. Mutant *unc79*^F01615^ (red) is less active than w1118 (grey) (N≥32; P<0.0001), while *unc79*^F03453^ (maroon) (N=32, P>0.0648), and heterozygous controls (pink) (N=32; P>0.0733). All sleep data are violin plots and SR data are survival curves. ***p < 0.001; ****p < 0.0001.

**Supplemental Figure 3.**

**A**. Knockdown of unc79 in different brain regions show no differences to respective flies expressing *luc*^RNAi^ (One-way ANOVA : F (12, 454) equal 2.247). Peptidergic knockdown of unc79 (C929>*unc79*^RNAi^) does not differ in control (C929> luc^RNAi^) (pink, P-value equals 0.9665, N>40). Fan-shaped body knockdown of unc79 (23E10>unc79 RNAi) does not differ in control (23E10>*luc*^RNAi^) (orange, P-value equals 0.698, N>29). Central complex knockdown of *unc79* (R69F08>*unc79*^RNAi^) does on differ in control (R69F08>*luc*^RNAi^) (green, P-value equals 0.8795, N>33). Glial knockdown of unc79 (repo>*unc79*^RNAi^)

(teal, P-value equals 0.9999, N>21) does on differ in control (Repo/luc^RNAi^). Circadian genes knockdown of unc79 timeless (tim> unc79 RNAi) (blue, P-value equals 0.8226, N>32) does on differ in control (tim/luc RNAi) and pigment dispersion factor (pdf> *unc79*^RNAi^) (purple, P-value equals 0.9997, N>32) does on differ in control (pdf/luc RNAi).

**B**. Starvation resistance for knockdown of *unc79* in brain region drivers compared to *luc* control (Gehan-Breslow-Wilcoxon test). Peptidergic knockdown of unc79 (C929>*unc79*^RNAi^) does not differ in control (C929/luc RNAi) (pink, df equal 1, P-value equals 0.668, N>40). Fan-shaped body knockdown of unc79 (23E10>unc79 RNAi) does not differ in control (23E10/luc^RNAi^) (orange, df equals 1, P-value equals 0.0879, N>32). Central complex knockdown of unc79 (R69F08>*unc79*^RNAi^) does on differ in control (R69F08>*luc*^RNAi^) (green, df equals 1, P-value equals 0.6268, N>33). Glial knockdown of unc79 (repo>*unc79*^RNAi^) (teal, df equals 1, P-value equals 0.3464, N>21) does on differ in control (Repo>*luc*^RNAi^). Circadian genes knockdown of unc79 timeless (tim>*unc79*^RNAi^) (blue, df equals 1, P-value equals 0.072, N>32) does on differ in control (tim>luc^RNAi^). Pigment dispersion factor (pdf>*unc79*^RNAi^) (purple, df equals 1, P-value equals 0.0191, N>32) is significantly different than control (pdf>*luc*^RNAi^). All sleep data are violin plots and SR data are survival curves. ***p < 0.001; ****p < 0.0001.

**Supplemental Figure 4.**

Table depicting average circadian metrics under free running conditions.

